# Tuning the sensitivity of genetically encoded fluorescent potassium indicators through structure-guided and genome mining strategies

**DOI:** 10.1101/2021.10.07.463355

**Authors:** Cristina C. Torres Cabán, Minghan Yang, Cuixin Lai, Lina Yang, Fedor Subach, Brian O. Smith, Kiryl D. Piatkevich, Edward S. Boyden

**Affiliations:** McGovern Institute for Brain Research, MIT, Cambridge, MA, USA; Department of Biological Engineering, MIT, Cambridge, MA, USA; Department of Media Arts & Sciences, MIT, Cambridge, MA, USA; School of Life Sciences, Westlake University, Hangzhou, Zhejiang Province, China; Westlake Laboratory of Life Sciences and Biomedicine, Hangzhou, Zhejiang Province, China; Institute of Basic Medical Sciences, Westlake Institute for Advanced Study, Hangzhou, Zhejiang Province, China; College of Physics, Jilin University, Changchun, Jilin Province, China; Complex of NBICS Technologies, National Research Center “Kurchatov Institute”, Moscow, Russia; Institute of Molecular, Cell & Systems Biology, College of Medical Veterinary & Life Sciences, University of Glasgow, Glasgow, UK; Koch Institute for Integrative Cancer Research, MIT, Cambridge, MA, USA; Howard Hughes Medical Institute, Chevy Chase, MD, USA; Department of Brain and Cognitive Sciences, MIT, Cambridge, MA, USA; K. Lisa Yang Center for Bionics, MIT, Cambridge, MA, USA; Center for Neurobiological Engineering, MIT, Cambridge, MA, USA

## Abstract

Genetically encoded potassium indicators lack optimal binding affinity for monitoring intracellular dynamics in mammalian cells. Through structure-guided design and genome mining of potassium binding proteins, we developed green fluorescent potassium indicators with a broad range of binding affinities. KRaION1, based on the insertion of a potassium binding protein (Ec-Kbp) into the fluorescent protein mNeonGreen, exhibits an isotonically measured K_d_ of 69±10 (mM; mean ± standard deviation used throughout). We identified Ec-Kbp’s binding site using NMR spectroscopy to detect protein-thallium scalar couplings and refined the structure of Ec-Kbp in its potassium-bound state. Guided by this structure, we modified KRaION1, yielding KRaION2, which exhibits an isotonically measured K_d_ of 96±9 (mM). We identified four Ec-Kbp homologs as potassium binding proteins, which yielded indicators with isotonically measured binding affinities in the 39-112 (mM) range. KRaIONs expressed and functioned in HeLa cells, but exhibited lower K_d_ values, which were mirrored by lower K_d_ values measured in vitro when holding sodium constant. Thus, potassium indicator K_d_ may need to be evaluated in the context of a given experimental goal.

## Introduction

Potassium ions are important for physiological functions including serving key roles in multipurpose electrochemical gradients of cells. Potassium dynamics have been studied in a range of cells and systems, including neurons and glia (Newman *et al*, 1984; Gardner-Medwin & Nicholson, 1983; Ficker & Heinemann, 1992; Mitterdorfer & Bean, 2002; Martina *et al*, 2007; Stansfeld *et al*, 1986), cardiomyocytes (Ocorr *et al*, 2007), and renal cells (Luo *et al*, 2016; Nie *et al*, 2005), and dysregulation of potassium homeostasis has been proposed in the manifestation of pathological conditions including seizures, immune cell impairment against cancer, and ischemic events (Eil *et al*, 2016; Windmüller *et al*, 2005; Barcia *et al*, 2012). Historically, potassium concentration in cells, tissues, and organisms has been measured with the use of electrodes or synthetic dyes (Octeau *et al*, 2018; Meeks & Mennerick, 2007; Bossy-Wetzel *et al*, 2004). However, there is still a lack of widely used methods to measure potassium in biological systems in a non-invasive and cell-targetable manner, as can be done with genetically encoded fluorescent indicators (Chen *et al*, 2013).

Recently the potassium binding protein, Kbp, cloned from *E. coli* (which we call *E. coli*-Kbp or Ec-Kbp for short), was reported to undergo large conformational changes during potassium binding. Ec-Kbp consists of ∼150 amino acids forming two domains: LysM (short for lysin motif) and BON (bacterial OsmY and nodulation). It binds to a single potassium ion, however the exact potassium binding site was not previously identified when it was first characterized (Ashraf *et al*, 2016). Upon binding to a potassium ion, Ec-Kbp undergoes a conformational change that brings the two domains closer together and orders the N-terminus such that it lies close to the C-terminus. This behavior parallels that of calmodulin and M13, where calmodulin’s two EF-hand domains change conformation upon calcium binding and capture the M13 peptide, bringing the N- and C-termini of the fusion closer together to cause fluorescence emission changes in commonly used genetically encoded calcium indicators. Ec-Kbp’s conformational change thus makes it a good candidate module for incorporation into genetically encoded sensors, particularly with the use of split-fluorescent proteins, as is used in the design of the bright green fluorescent calcium indicator NCaMP7 (Subach *et al*, 2020).

Immediately following the identification of Ec-Kbp as a potassium binding protein, the protein engineering community started using it to develop genetically encoded K^+^ sensors. To date, two FRET sensors, GEPII 1.0 and KIRIN1, which comprise fusion proteins of a donor-acceptor pair of fluorescent proteins fused to the termini of Ec-Kbp, and whose fluorescence emission properties change depending on the binding of K^+^, have been published (Bischof *et al*, 2017; Shen *et al*, 2019). A single-fluorescent protein K^+^ sensor, GINKO1, has also been developed by inserting Ec-Kbp into EGFP to modulate its fluorescence in a K^+^ concentration dependent manner (Shen *et al*, 2019). Although these sensors have been used to measure K^+^ concentration in solution and in cell culture, their binding affinities for K^+^ are not appropriate for accurate measurements of potassium concentration inside cells (previously reported K_d_ values range from 0.40-2.6 (mM)), whereas textbook measurements of K^+^ concentrations lie in the range from 140-150 mM for mammalian cells at resting potential (Somjen, 1979; Thier, 1986). Extracellular K^+^ concentrations can fluctuate up to 8-12 mM during processes such as prolonged neuronal firing, and can fluctuate even more (by 30-80 mM) in events of sustained depolarization such as spreading depression (Heinemann & Lux, 1977; Futamachi *et al*, 1974; Lothman *et al*, 1975). The excess potassium can be taken up and redistributed by glial cells, which can see an increase of up to ∼60% in intracellular K^+^ concentration during such events (Dufour *et al*, 2011; Amzica *et al*, 2002). However, without the ability to measure K^+^ concentration dynamically in a cell-specific way, we will not know what K^+^ dynamics look like in the vast majority of cell types of the body in different healthy and disease states. Thus, our goal was to design and generate intracellularly expressed, genetically encoded potassium sensors, with an initial focus on developing a sensor with a K_d_ value that makes it sensitive to changes in K^+^ comparable to those in the intracellular milieu.

We here report our progress towards this goal, resulting in a family of genetically encoded green fluorescent sensor prototypes generated via two approaches. One approach was guided by the NMR characterization of Ec-Kbp’s potassium binding site while the other was driven by identifying and utilizing a related set of alternative potassium binding proteins. In each case, we fused the mammalian codon-optimized gene to that encoding for mNeonGreen. With the former approach, we used structural information to guide mutagenesis of Ec-Kbp’s binding site to tune its K_d_. Two of the mutants generated exhibited binding affinity to K^+^, measured under isotonic conditions, in the range of 96-138 (mM) and fluorescence dynamic range of ∼200% when increasing K^+^ concentrations from 0.1 to 150 mM under isotonic conditions. With the latter approach, we identified previously unannotated proteins from metagenomic sequencing databases that share 45-72 % amino acid identity with Ec-Kbp. We incorporated these proteins as the potassium sensing moiety in the indicator, and discovered that the resulting indicators display isotonically measured potassium binding affinities ranging from 39-112 (mM) and a fluorescence dynamic range from ∼100-200 % over the range from 0.1-150 mM K^+^. All of the generated indicators can be excited by two wavelengths, which enables ratiometric imaging, which in turn mitigates artifacts during imaging, for example accounting for variable indicator concentrations, and compensating for photobleaching of the fluorophore, changes in laser intensity, and other standard aspects of fluorescence imaging. In summary, we demonstrate an approach to K^+^ sensor protein design by which we generate a set of single fluorescent protein K^+^ indicators that display a wide range of isotonically measured binding affinities to potassium in vitro. However, when expressed in HeLa cells, all indicators tested had K_d_ values at a lower range in comparison to the measurements obtained in vitro under isotonic measurement conditions. And, when we measured K_d_ in vitro under different conditions, holding sodium constant while varying potassium, we obtained generally lower K_d_ values than observed under isotonic conditions. Thus, K_d_ could vary depending on the conditions used for its measurement, and perhaps, in the live cell case, unwanted interactions with other intracellular components (Rana *et al*, 2019). We thus consider these indicators to remain in the prototype stage, as discrepancies in binding affinities of the indicators may require thoughtful consideration of how K_d_ is measured in cellular and environmental contexts, for the case of potassium. Future work will be needed to identify the proper calibration conditions in cell types of interest, and applied to the indicators described here, and elsewhere.

## Results

### Development of a genetically encoded green potassium sensor

We sought to design a genetically encoded potassium indicator that would utilize a single fluorescent protein as the reporting moiety. Previously developed potassium indicators have either used a FRET design, which requires the use of two fluorescent fusion partners (Shen *et al*, 2019; Bischof *et al*, 2017), or have used direct insertion of Ec-Kbp into EGFP (Shen *et al*, 2019). We decided to utilize an approach similar to the latter design, that is based on the insertion of the sensing moiety (in this case, the potassium binding moiety) into a fluorescent protein as has been done with other indicators for calcium: ncpGCaMP6s, NTnC and NCaMP7 (Qian *et al*, 2019; Barykina *et al*, 2016; Subach *et al*, 2020). Direct insertion of the sensing moiety into a fluorescent protein is an alternative approach to circular permutation of the fluorescent protein or sensing moiety (Nasu *et al*, 2021). Of the sensors that utilize this design, NCaMP7, which contains mNeonGreen (mNG) as a green fluorescent reporter, is 1.7-fold brighter than another commonly used calcium indicator GCaMP6s (Chen *et al*, 2013) and has a maximum ΔF/F = 2700%. Our potassium indicator design thus used NCaMP7 as a template, as mNeonGreen is brighter than other characterized monomeric GFP proteins (Shaner *et al*, 2013), and the sensor design allows for direct insertion of the binding domain into the fluorescent protein.

Relative to NCaMP7, we replaced the calmodulin/calmodulin binding peptide module with the potassium binding protein, Ec-Kbp, while preserving the original amino acid linkers between the fluorescent protein and the binding domain (**Figure 1a, b**). Assessment of the obtained construct mNG-Ec-Kbp in crude bacterial extract revealed significant changes in green fluorescence intensity upon increased potassium ion concentration at physiological pH. Encouraged by the evident potassium responsiveness, we purified mNG-Ec-Kbp and performed detailed characterization of its spectroscopic and biochemical properties at 0 and 150 mM under isotonic conditions (**Table 1**). Since the spectroscopic characterization revealed a ratiometric fluorescence response by excitation, we named this sensor KRaION1, which stands for <u>K</u>^+^ <u>ra</u>tiometric indicator for <u>o</u>ptical imaging based on m<u>N</u>eonGreen. The absorbance spectrum of KRaION1 shows two major bands with peaks at 408 and 508 nm (**Figure 1c**), which in mNG corresponds to the protonated-neutral and deprotonated-anionic forms of the chromophore, respectively (Steiert *et al*, 2018). Upon potassium administration, the peak at 508 nm increased while the peak at 408 nm decreased. Correspondingly, the excitation fluorescence spectrum of KRaION1 has two bands with peaks at 407 nm and 507 nm (**Figure 1d**). Excitation of either peak resulted in green fluorescence with identical fluorescence emission maximum at 518 nm. The indicator’s extinction coefficient (EC) ratio was 65,300/12,000 M^-1^cm^- 1^ when excited at 408/507 at 0 mM K^+^ and 45,600/23,200 M^-1^cm^-1^ when excited at 407/507 at 150 mM K^+^. The corresponding quantum yields obtained at 0 mM and 150 mM K^+^ when excited at 407/507 nm were 35/26% and 29/57%, respectively (**Table 1**). Therefore, upon potassium administration the green fluorescence excited at 507 nm increases due to both increasing QY and EC, while fluorescence excited at 407 nm decreases due to both decreasing QY and EC. This confirmed that KRaION1 is a ratiometric indicator, which can be excited by two wavelengths and can be read out by the same emission wavelength.

**Figure 1.**
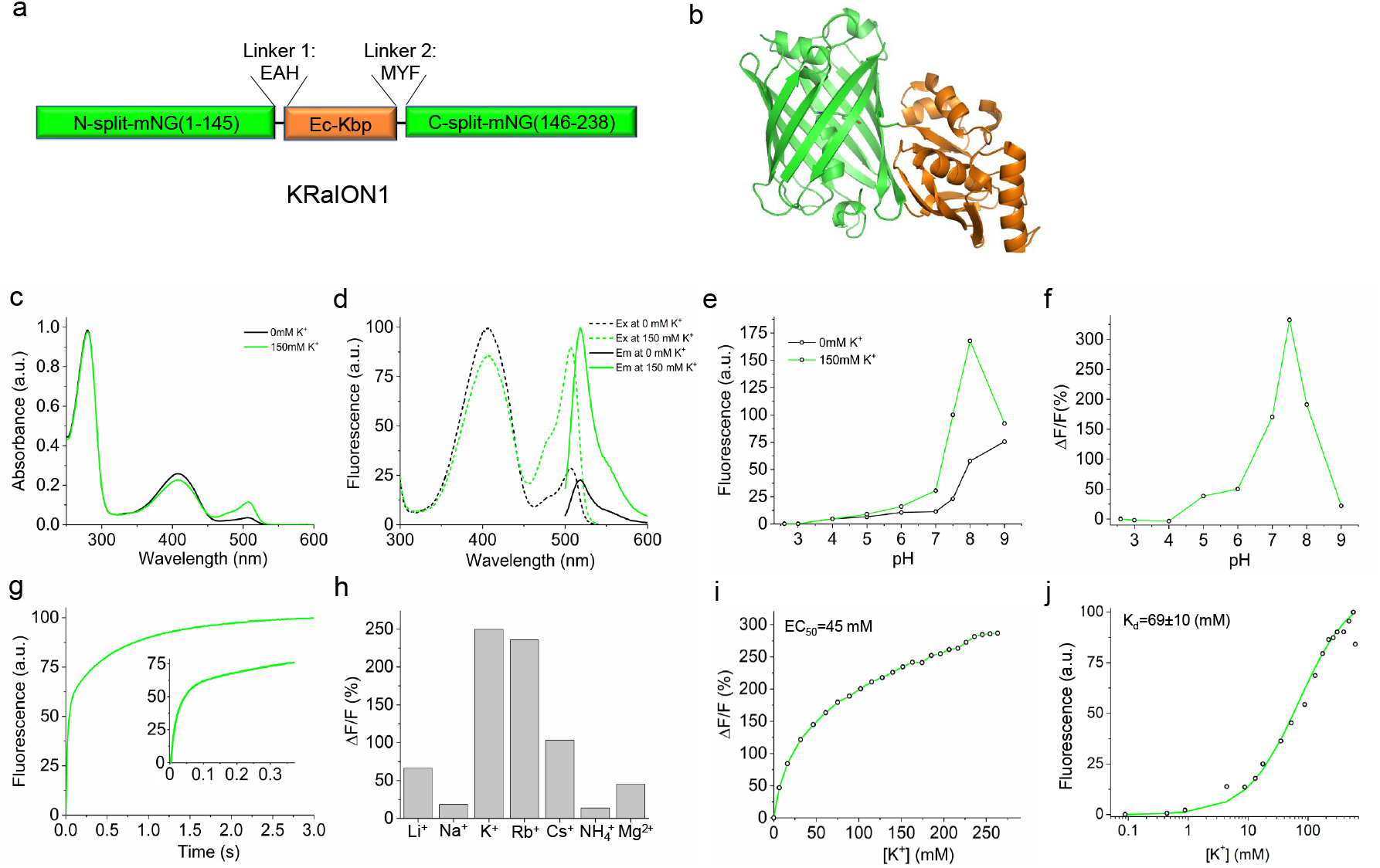
Molecular design and properties of the genetically encoded green potassium indicator KRaION1 in solution. (a) Molecular design of the KRaION1 indicator. (b) The proposed structure of KRaION1 shown as a ribbon diagram according to the crystal structure of NCaMP7 (PDB:6XW2) and NMR structure of the Ec-Kbp domain (PDB:5FIM). (c) Absorbance spectra of KRaION1 at 0 mM and 150 mM potassium at pH = 7.4. (d) Fluorescence spectra of KRaION1 at 0 mM and 150 mM potassium at pH = 7.4. (e) Relative fluorescence intensity of KRaION1 at 0 mM and 150 mM potassium as a function of pH. (f) Fluorescence changes upon addition of 150 mM potassium as a function of pH at constant ionic strength. (g) Potassium-association using stopped-flow fluorimetry. Association kinetics curves were acquired at 40 mM final K^+^ concentration starting from K-free protein solution. Small graph is the same association kinetics curve shown in the range of 0 - 300 ms timeframe. (h) Selectivity of KRaION1 to K^+^ and other cations measured by titration of Li^+^, Na^+^, Rb^+^, Cs^+^, NH_4_^+^ and Mg_2_^+^. Calculated ΔF/F % for each cation was measured in the range of 0 - 260 mM. (i) Fluorescent dynamic range and EC_50_ of KRaION1 measured in the range of 0 – 260 mM K^+^. (j) Potassium titration data points (open circles) measured at pH = 7.3 and constant ionic strength of 700 mM, fitted using Q = (Q_max_ - Q_0_)Y+Q_0_ function (green line).

**Table 1.**
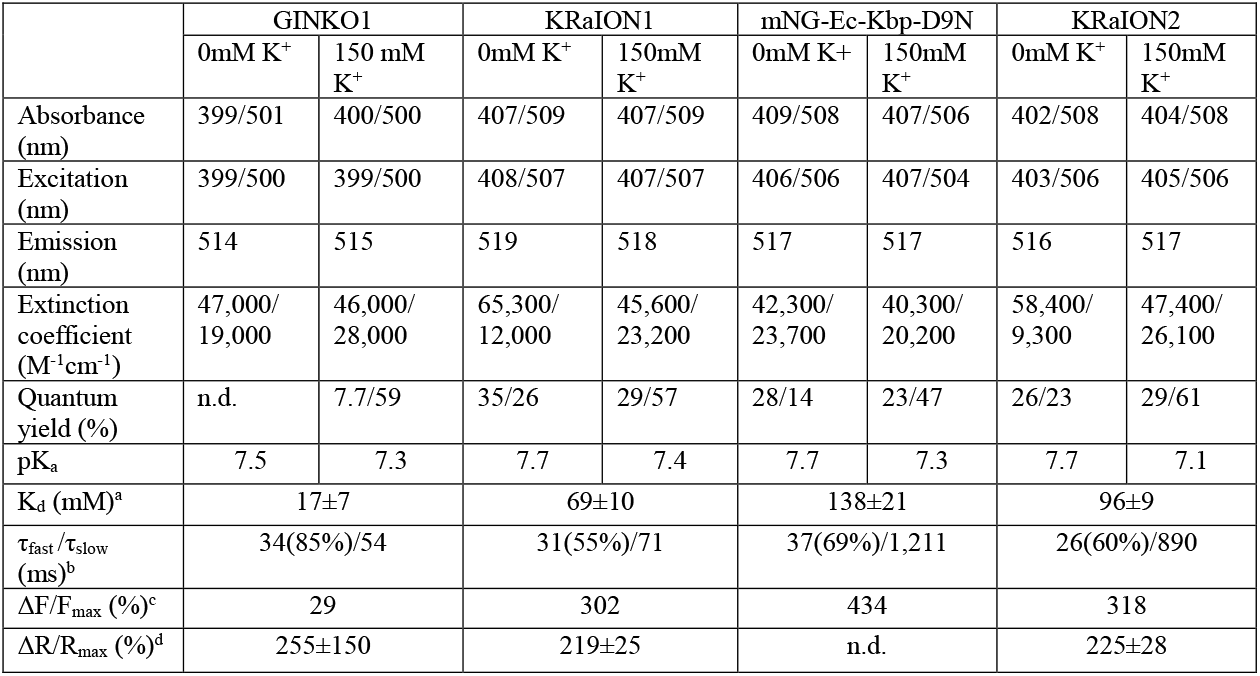
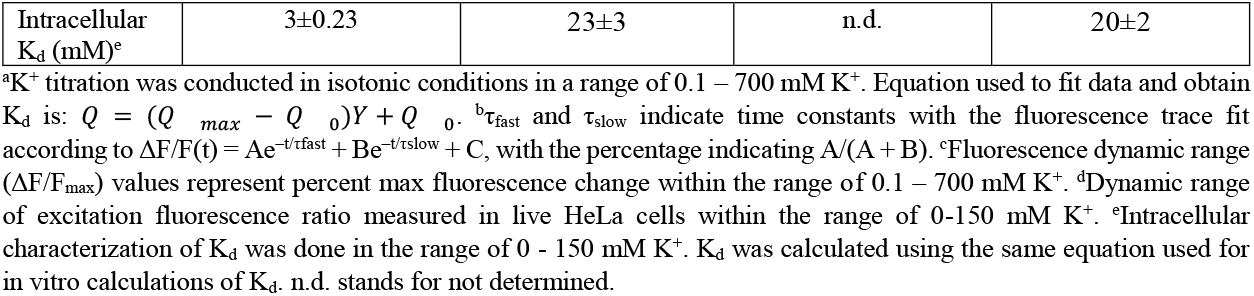
Spectral and biochemical properties of green fluorescence potassium sensors in solution.

|  | GINKO1 |  | KRaION1 |  | mNG-Ec-Kbp-D9N |  | KRaION2 |  |
| --- | --- | --- | --- | --- | --- | --- | --- | --- |
| | 0mM $\text{K}^+$ | 150 mM $\text{K}^+$ | 0mM $\text{K}^+$ | 150mM $\text{K}^+$ | 0mM $\text{K}^+$ | 150mM $\text{K}^+$ | 0mM $\text{K}^+$ | 150mM $\text{K}^+$ |
| Absorbance (nm) | 399/501 | 400/500 | 407/509 | 407/509 | 409/508 | 407/506 | 402/508 | 404/508 |
| Excitation (nm) | 399/500 | 399/500 | 408/507 | 407/507 | 406/506 | 407/504 | 403/506 | 405/506 |
| Emission (nm) | 514 | 515 | 519 | 518 | 517 | 517 | 516 | 517 |
| Extinction coefficient ($\text{M}^{-1}\text{cm}^{-1}$) | 47,000/19,000 | 46,000/28,000 | 65,300/12,000 | 45,600/23,200 | 42,300/23,700 | 40,300/20,200 | 58,400/9,300 | 47,400/26,100 |
| Quantum yield (%) | n.d. | 7.7/59 | 35/26 | 29/57 | 28/14 | 23/47 | 26/23 | 29/61 |
| $\text{pK}_a$ | 7.5 | 7.3 | 7.7 | 7.4 | 7.7 | 7.3 | 7.7 | 7.1 |
| $K_d$ (mM)<sup>a</sup> | 17±7 | | 69±10 | | 138±21 | | 96±9 | |
| $\tau_{\text{fast}}/\tau_{\text{slow}}$ (ms)<sup>b</sup> | 34(85%)/54 | | 31(55%)/71 | | 37(69%)/1,211 | | 26(60%)/890 | |
| $\Delta F/F_{\text{max}}$ (%)<sup>c</sup> | 29 | | 302 | | 434 | | 318 | |
| $\Delta R/R_{\text{max}}$ (%)<sup>d</sup> | 255±150 | | 219±25 | | n.d. | | 225±28 | |

|  |  |  |  |  |
| --- | --- | --- | --- | --- |
| Intracellular<br>K<sub>d</sub> (mM)<sup>e</sup> | 3±0.23 | 23±3 | n.d. | 20±2 |

<sup>a</sup>K<sup>+</sup> titration was conducted in isotonic conditions in a range of 0.1 – 700 mM K<sup>+</sup>. Equation used to fit data and obtain K<sub>d</sub> is: $Q = (Q_{max} - Q_0)Y + Q_0$ . <sup>b</sup> $\tau_{fast}$ and $\tau_{slow}$ indicate time constants with the fluorescence trace fit according to $\Delta F/F(t) = Ae^{-t/\tau_{fast}} + Be^{-t/\tau_{slow}} + C$ , with the percentage indicating A/(A + B). <sup>c</sup>Fluorescence dynamic range ( $\Delta F/F_{max}$ ) values represent percent max fluorescence change within the range of 0.1 – 700 mM K<sup>+</sup>. <sup>d</sup>Dynamic range of excitation fluorescence ratio measured in live HeLa cells within the range of 0-150 mM K<sup>+</sup>. <sup>e</sup>Intracellular characterization of K<sub>d</sub> was done in the range of 0 - 150 mM K<sup>+</sup>. K<sub>d</sub> was calculated using the same equation used for in vitro calculations of K<sub>d</sub>. n.d. stands for not determined.

The indicator’s functionality was tested at different pH concentrations to determine its pH sensitivity. The KRaION1 indicator exhibits a large fluorescence change in response to 150 mM K^+^ within the physiologically relevant pH range of 7 – 8. The maximum dynamic range (ΔF/F) of 332% was seen at pH 7.5 (**Figure 1e, f**). At pH levels outside this range, the fluorescence response of the indicator decreased to less than 50%. Taken together, these data suggest that KRaION1 would be sensitive to fluctuations in pH, which is typical for the majority of other single fluorescent protein-based green indicators (Qian *et al*, 2019; Barykina *et al*, 2016; Subach *et al*, 2020). Monitored in solution at pH = 7.4, KRaION1 displayed a fluorescence response upon binding K^+^ (at final [K^+^] = 40 mM) that was bi-exponential with time constants of τ_fast_ = 31 ms, which accounts for 55% of the total amplitude, and τ_slow_ = 71 ms, that accounts for the remaining amplitude (**Figure 1g**). This association kinetics response was comparable to that for GINKO1 under identical conditions, characterized by τ_fast_ = 34 ms, which accounts for 85% of its total amplitude, and τ_slow_ = 54 ms that accounts for the remaining amplitude. The fluorescence response of KRaION1 is about 3-fold faster compared to NCaMP7, which perhaps could be due to the faster association kinetics of potassium ions with Ec-Kbp. The association kinetics of KRaION1 could be suitable to measure some of the fast potassium currents observed in neurons and glia (Luther *et al*, 2000; Sibille *et al*, 2015).

Since the cellular environment contains several metal ions, including sodium and magnesium, it is important to measure KRaION1’s ion selectivity. We measured fluorescence changes upon titrations with the following ions: Li^+^, Na^+^, Rb^+^, Cs^+^, NH_4_^+^, and Mg^2+^, along with K^+^ for comparison (**Figure 1h**). There were small, or almost no, changes in fluorescence with all ions except for Rb^+^ and Cs^+^, consistent with what was observed in the initial characterization of Ec-Kbp (Ashraf *et al*, 2016). Next, we determined the fluorescence dynamic range of KRaION1 under conditions mimicking intracellular ion compositions (Lodish *et al*, 2000). For this, we measured fluorescence changes of KRaION1 with ∼10 mM steps in potassium concentration until the fluorescence reached plateau. The green fluorescence increased to about 286% of ΔF/F at ∼250 mM K^+^. The corresponding EC_50_ value was approximately 45 mM (**Figure 1i**).

The isotonically characterized binding affinity or K_d_ of KRaION1 was obtained as 69±10×10^−3^ (69±10 (mM)) by fitting the measured green fluorescence response of the sensor at different K^+^ concentrations under isotonic conditions to a single site binding model (**Figure 1j**). For comparison, the EGFP-based potassium indicator, GINKO1, under similar conditions had a K_d_ value of 17±7×10^−3^ (17±7 (mM)). These K_d_ values, although useful in other biological contexts, would be less useful in detecting small changes in mammalian cells’ intracellular potassium concentration (∼150 mM K^+^). We therefore sought to optimize the prototype sensor KRaION1 to achieve a higher K_d_ that would provide better sensitivity to small changes in potassium concentration under intracellular conditions in mammalian cells. To achieve this aim we sought additional information about Ec-Kbp’s ion binding site.

### Identification of Ec-Kbp’s ion binding site

Since Ec-Kbp has so far proved refractory to crystallization, we used NMR spectroscopy to locate Ec-Kbp’s ion binding site. Isotopes of potassium are quadrupolar nuclei with modest sensitivity making them unsuitable for determining protein-ion contacts in solution. Instead, we used thallium(I) as a K^+^ mimic allowing the measurement of direct two bond scalar couplings (^2^J ^13^C=O…^205/203^Tl^+^) (B.O. Smith, in preparation) of around 100 Hz in the HNCO spectrum (**Figure 2a**) that unambiguously identify the backbone carbonyls of V7 and A10 from loop 1 in Ec-Kbp’s N-terminal extension and G75, I77 and I80 from loop 5, the turn immediately preceding the BON domain’s last β-strand, as Tl^+^ ligands.

**Figure 2.**
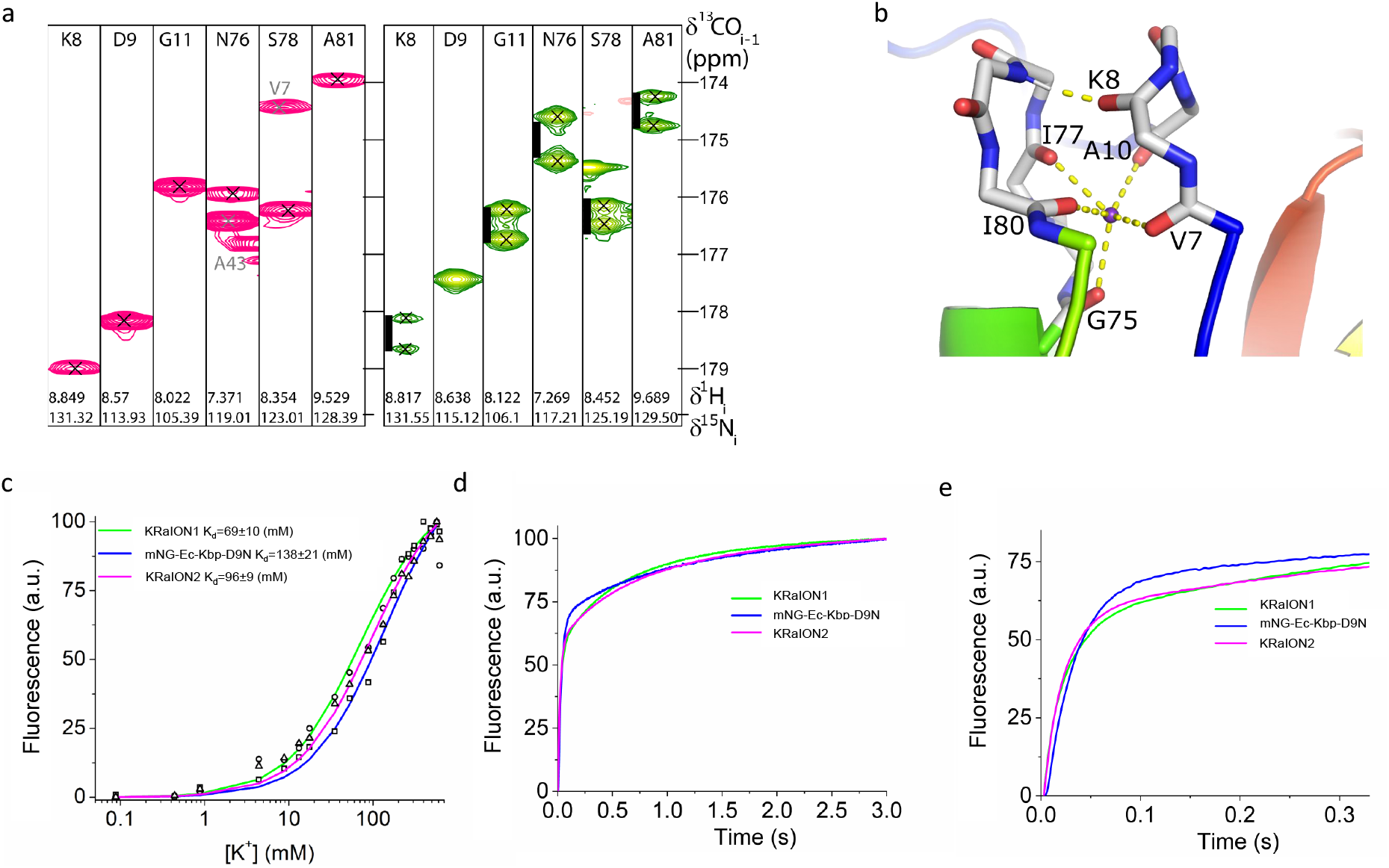
Identification and modification of Ec-Kbp’s potassium binding site. (a) ^1^H,^13^C strips from HNCO spectra of K^+^ (magenta) and Tl^+^ (green) bound Ec-Kbp showing the residues whose crosspeaks are split by an additional J-coupling in the Tl+ bound form. The black scale bar represents 100 Hz. (b) Refined configuration of the Ec-Kbp ion binding site. The ion is coordinated entirely by backbone carbonyls from loops 1 & 5 (backbone atoms shown as sticks). Sections of the rainbow-colored cartoon of the rest of the protein are visible. Coordinating carbonyl to K^+^ interactions and the K8 to G79 hydrogen bond are shown as yellow dashed lines. (c) Potassium titration data points for KRaION1 and mutants mNG-Ec-Kbp-D9N and KRaION2 (open circles, squares, and triangles, respectively) measured at pH = 7.4 and constant ionic strength, fitted using the equation Q = (Q_max_ - Q_0_)Y+Q_0_ (denoted by a green line, blue line, and magenta line, respectively). (d) Potassium-association using stopped-flow fluorimetry. Association kinetics curves were acquired at 40 mM final K^+^ concentration starting from K-free protein solution. (e) Association kinetics curves in the range of 0 – 300 ms timeframe.

### Refined structure of Ec-Kbp.K^+^

To provide an accurate structural model on which to base mutagenesis of the potassium binding domain of KRaION1, we recalculated the Ec-Kbp.K^+^ structure (Ashraf *et al*, 2016) incorporating a K^+^ ion restrained to the carbonyl oxygens identified as Tl^+^ ligands (PDB:7PVC; **Supplementary Table 1**). The coordinating carbonyl oxygens were restrained to be between 2.75±0.1 Å of the K^+^ ion, as suggested by Harding (Harding, 2006) without any constraint on the C=O…K^+^ bond angle. Since the NMR structure calculation software, ARIA2.3/CNS, uses the OPLX forcefield with explicit TIP3P-like water representation for the water refinement stage (Linge *et al*, 2003) we used the parameters for K^+^ suggested by Mamatkulov & Schwierz (Mamatkulov & Schwierz, 2018). The refined structure ensemble differs little from the structure calculated without an ion included (**Supplementary Figure 1a**) but binds K^+^ between loops 1 and 5 at the centre of a distorted square pyramid whose base of approximately 3.3-4.7 on each side is formed by the carbonyl oxygens of residues V7, A10, I77 and I80, while the carbonyl oxygen of residue G75 sits at the apex with the lateral sides ranging from 3.56±0.40 for G75 and I80 to 5.05±0.29 for G75 and V7. Each of the five backbone carbonyls coordinate the K^+^ at distances between 2.62 and 2.86 (**Table 2**). The carbonyl oxygen of K8 appears a candidate as a sixth K^+^ ligand forming the vertex opposite G75’s carbonyl oxygen in an elongated octahedral coordination sphere. However, K8’s carbonyl oxygen lies more distant from the K^+^ (3.98±0.37 Å) and is instead involved in a hydrogen bond with the backbone amide of G79 (**Figure 2b**). Each of the conserved glycine residues in the two loops performs a critical role in allowing the ion binding site to form. G11 adopts an extended conformation with a positive φ dihedral angle and its lack of sidechain allowing the close approach of the two loops. G75 is in an alpha-helical conformation, but the sidechain of any other residue at this position would disrupt the close packing of the conserved F6 with its neighboring residues. G79 contributes to a tight turn with a positive φ allowing its amide to hydrogen bond with K8 as noted above. The other conserved residue in loop5, N76, has a critical role in the overall fold of the protein as its side-chain amide makes a pair of inter-domain hydrogen bonds with V143 in the LysM domain. As well as contributing directly to ion binding with its backbone carbonyl, residue 10 is conserved as an alanine, likely to ensure the correct close contact can be made both with V7 and with G141 at the N-terminus of the final beta-strand in the LysM domain.

**Table 2.**
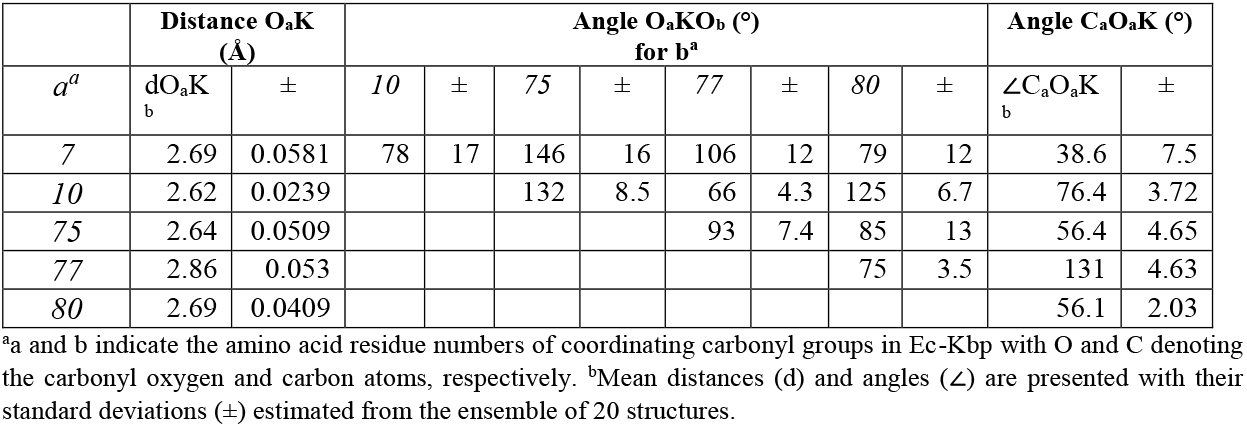
Geometry of the K^+^ binding site in Ec-Kbp.K.

### Structure-guided mutagenesis modulates K^+^ binding

In order to fine-tune the potassium binding affinity of the Ec-Kpb domain, we decided to introduce mutations within and around the ion binding site, informed by the structure determined by NMR (**Figure 2b** and **Supplementary Figure 1b**). Since the K^+^ ion appears to be coordinated exclusively by the protein’s backbone carbonyl groups, our scope for directly affecting the binding site was limited. Instead, we speculated that adjusting the electrostatic character of residues near the binding pocket could affect the stability of the K^+^ bound state and shift the K_d_ to higher values that lie within the physiological range of intracellular K^+^ concentration that is found in mammalian cells. To test this hypothesis, we generated a set of KRaION1 mutants with altered residues within and near the K^+^ binding site. The binding site residues K8, A10, and N76 were swapped for another small residue (A, G), or to a negatively charged group D. These choices were made based on the electrostatic characteristics of the residues: positive or neutral residues were swapped for other neutral or negative residues, respectively. The conserved hydrophobic residues I77 and I80 were also mutated to smaller amino acids that are either neutral, such as A and G, or small and polar, such as S. Guided by the structure of Ec-Kbp, we also deduced that the nearby residues D9 and E12 could be mutated to remove their negative charges and thus make the local electrostatic environment less favorable for K^+^ binding without disrupting binding altogether. These residues were substituted with either polar, positively charged, or small non-polar residue groups: N/A/K or Q/A/K, respectively.

We purified each of the generated KRaION1 mutants and measured their fluorescence and absorbance spectra in solution over a range of K^+^ concentrations from 0 mM to 310 mM under non-isotonic conditions at pH = 7.3. We determined their properties including apparent K_d_, ΔF/F, baseline fluorescence brightness, and protein folding efficiency **(Supplementary Table 2**). We used non-isotonic conditions for quick screening of the generated variants’ functionality, but for proper assessment of the K_d_ of selected variants, we used isotonic conditions as we aimed to mimic the constant ionic strength maintained in the intracellular environment. It’s of note that KRaION1’s apparent K_d_ in these non-isotonic conditions was 12±4 (mM) in comparison to a K_d_ of 69±10 (mM) in isotonic conditions, which suggests the indicator’s binding affinity is affected by the ionic strength of the environment; thus, we proceeded with K_d_ measurements obtained in an environment more closely resembling that of cells. Of the KRaION1 mutant variants generated, mutations made directly to the K^+^ binding site appeared to disrupt proper functioning of the indicator, as we observed fluorescence dynamic ranges of < 30 % and folding efficiency of < 50 % in comparison to the original KRaION1. However, mutants D9N and E12A, which are adjacent to the K^+^ binding site, exhibited higher K_d_ values and fluorescence dynamic range in comparison to KRaION1. When characterized under the standard isotonic conditions mentioned earlier in the paper, D9N and E12A exhibited K_d_ values of 138±21×10^−3^ (138±21 (mM)) and 96±9×10^−3^ (96±9 (mM)), respectively (**Figure 2c**). This suggested that changing the electrostatic environment of the binding pocket is possible through rational design guided by the structure of the protein. Both mutants also exhibited different kinetics of association to K^+^ in comparison to KRaION1. Mutant D9N had time constants of activation of τ_fast_ = 37 ms accounting for 69% of total amplitude and τ_slow_ = 1,211 ms accounting for the remaining amplitude (**Figure 2d**). In contrast, mutant E12A had time constants of activation of τ_fast_ = 26 ms which accounted for 60% of total amplitude and τ_slow_ = 890 s which accounted for the remaining amplitude (**Figure 2e**). The D9N mutation led to slower activation kinetics, whereas the E12A mutation led to faster activation kinetics in relation to the initial indicator KRaION1. The mNG-Ec-Kbp-E12A mutant was thus named KRaION2 because of its desired K_d_ and kinetics values.

### Ec-Kbp homologs can be used as alternative binding moieties

As an alternative approach towards designing an indicator that had a more suitable binding affinity relevant to the intracellular mammalian environment, we searched for Ec-Kbp homologs that could be found in nature to use as the sensing moiety in our fluorescent biosensor. We used protein BLAST to search through a metagenomic dataset (env_nr), which contains proteins from whole genome shotgun sequencing (WGS) metagenomic projects. We selected four new small proteins with unknown function that shared 45-72% amino acid (aa) identity with the original Ec-Kbp and in which the K^+^ binding site was also largely conserved (**Supplementary Table 3, Figure 3a**). Protein homolog names are based on the organism or origin of sample collection. Ec: *E. coli* (GenBank: WP_000522415.1), C: compost (GenBank: MNG82101.1), Pa: *P. aeruginosa* (GenBank: NP_253865.1), Hv: hydrothermal vent (GenBank: VAV91021.1), D: *Defluviicoccus sp*. (GenBank: SUS08588.1). Importantly, any amino acid changes observed in the putative potassium binding site maintained the original residues’ electrochemical properties, supporting our choices of these proteins. Changes to binding site residue V7 to I7 were observed in D-Kbp. Changes to residue I77 to V at the equivalent positions were seen in all identified homologs, with the exception of Hv-Kbp whose residue change was to T. A final change was seen in the Pa-Kbp homolog, in which residue I80 was swapped for V.

**Figure 3.**
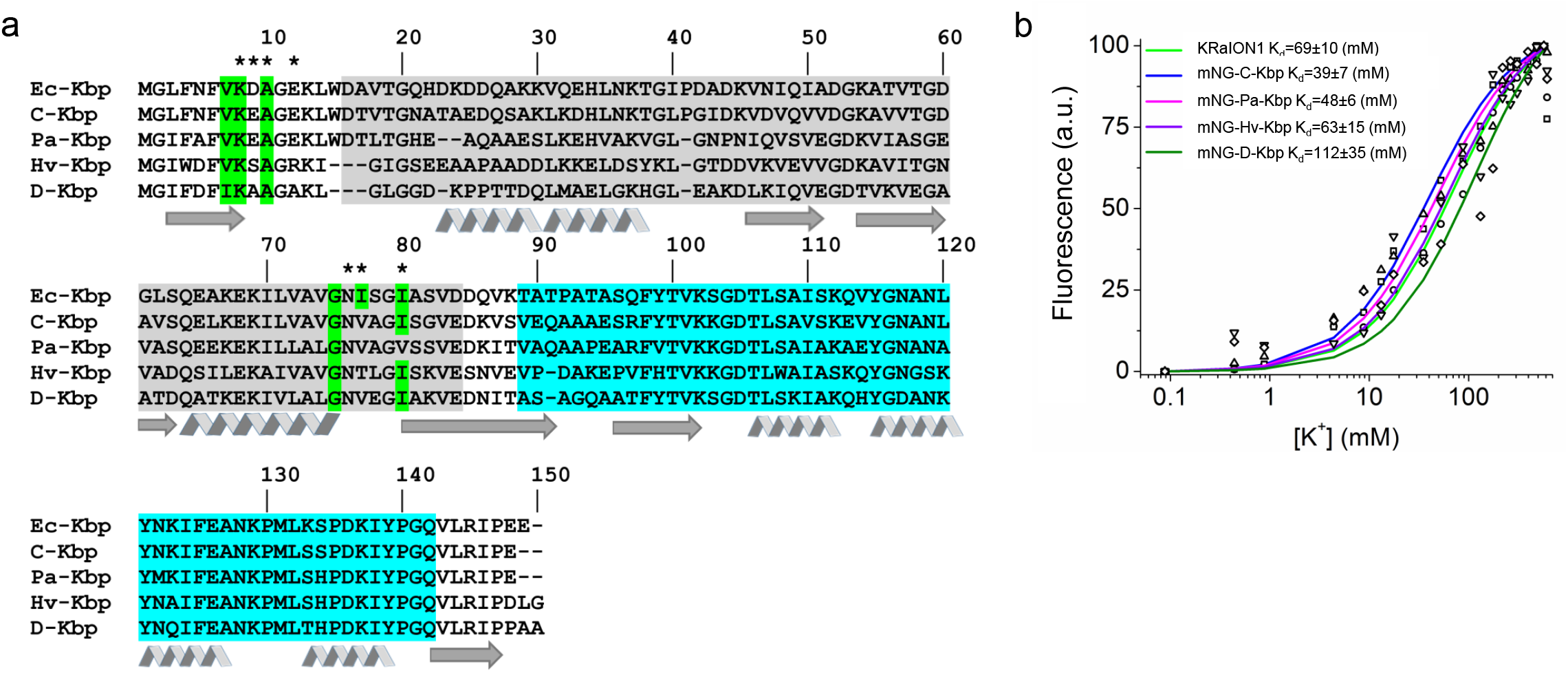
Identification of alternative potassium binding proteins and comparison to Ec-Kbp (a) Alignment of amino acid sequences of Ec-Kbp with five homologs obtained from a metagenomic BLAST search. Residues comprising the LysM and BON domains are shaded in grey and cyan, respectively. Residues highlighted in green indicate those that are conserved with the Ec-Kbp-identified potassium binding site. The β-sheet-forming regions and α-helix-forming regions are denoted by arrows and ribbons, respectively. Residues that were selected for site-directed mutagenesis in Ec-Kbp are denoted by asterisks. (b) Potassium titration data points for KRaION1 and homologs mNG-C-Kbp, mNG-Pa-Kbp, mNG-Hv-Kbp and mNG-D-Kbp (open circles, squares, upward triangles, downward triangles, and diamonds, respectively) measured at pH = 7.4 and constant ionic strength, fitted using the equation Q = (Q_max_ - Q_0_)Y+Q_0_ (denoted by a green line, blue line, magenta line, purple line, and dark green line, respectively).

We proceeded to exchange Ec-Kbp for each of these newly identified homologs in the KRaION1 indicator to check their functionality. As with the Ec-Kbp variants, we purified each of the four sensors designed with the homologs as alternative binding moieties and measured their fluorescence spectra at different K^+^ concentrations under isotonic conditions. They each exhibited a fluorescence response that varied with K^+^ concentration suggesting that these homologs are also K^+^ binding proteins and sequence changes near the binding site did not disable their K^+^ binding function (**Supplementary Table 4, Supplementary Figure 2**). These alternative binding moieties confer a range of isotonically characterized binding affinities to the indicators, from 39 (mM) to 112 (mM) (**Figure 3b**). One homolog, mNG-D-Kbp, displayed an isotonically measured K_d_ value of 112±35×10^−3^ (112±35 (mM)), in the same range as KRaION1 and its D9N and E12A variants. Thus, alternative potassium binding domains can generate promising potassium indicator candidates, focusing on the core parameter of K_d_ in the current study.

### In vivo characterization of indicators in HeLa cells

To characterize the response of the genetically encoded sensors inside mammalian cells, we focused on just a subset of these new variants, in particular, KRaION1 and KRaION2, in cultured HeLa cells. In addition to KRaION1, we chose the E12A mutant to test intracellularly as it has a combination of higher K_d_ and faster kinetics than the initial indicator. We excluded the D9N and mNG-D-Kbp variants from these experiments because of their slower kinetics and lower ΔF/F, respectively. We decided to take advantage of the ratiometric response of the sensors and used UV and cyan light to excite green fluorescence of both KRaION1 and KRaION2, which we found to be evenly distributed across the cytosolic space (**Figure 4a**). To permeabilize the cells and allow intracellular diffusion of potassium ions from outside the cell, we used the potassium ionophore valinomycin. We used a custom perfusion system to pump buffers with different concentrations of K^+^ into the extracellular space once the cells were permeabilized by valinomycin. In between buffer exchange, cells would be incubated in a buffer solution of 0 mM K^+^ to allow sensor equilibration to baseline fluorescence. The ratio of the fluorescence emission changes (F_487nm_/F_390nm_) showed KRaION2 responses when pulsed with K^+^ buffers ranging from 0 mM to 150 mM K^+^ (**Figure 4b**). This positive response suggests the indicators retain their function when expressed in cells. The ΔR/R_0_ increased with increasing K^+^ concentration, with the max ΔR/R_0_ percentage for KRaION2 intracellularly at 150 mM K^+^ as 225±28%. A similar max ΔR/R_0_ was obtained for KRaION1, 219±25%. The same experiment carried out in HeLa cells expressing GINKO1 showed a max ΔR/R_0_ of 255±150% (the significantly larger deviation of ΔR/R_0_ for GINKO1 across cells was due to its lower fluorescence brightness).

**Figure 4.**
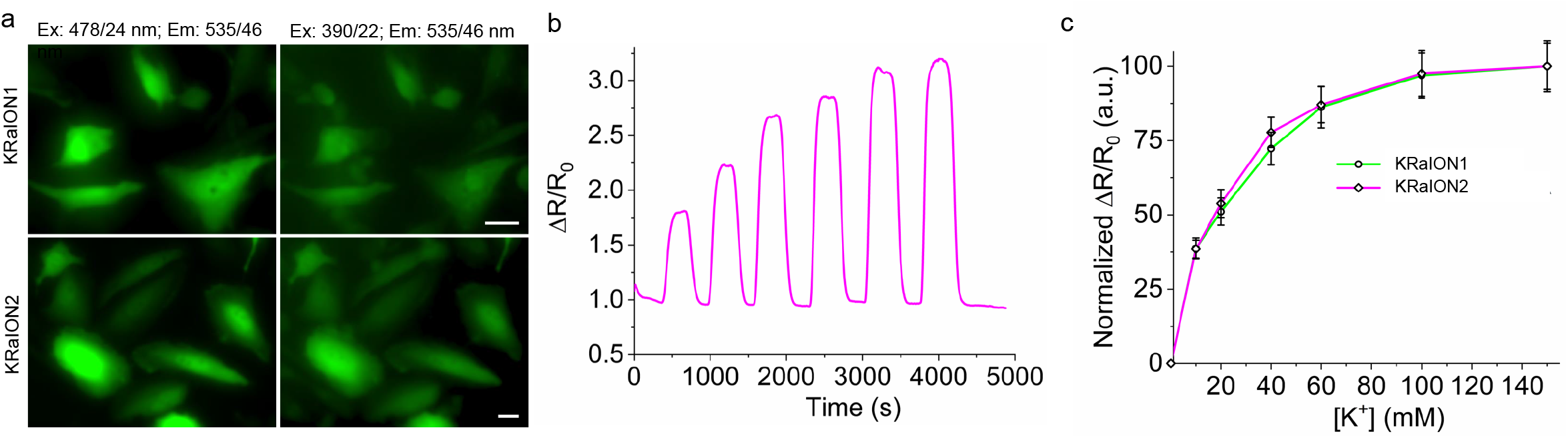
Calibration of KRaION1 and KRaION2 in live HeLa cells. (a) Representative fluorescence images of KRaION1 and KRaION2 in green channel with UV and cyan excitations (n=10 FOVs from three independent transfections each). Scale bars, 10 µm. (b) Representative excitation ratiometric response of KRaION2 at increasing [K^+^] from 0 to 150 mM (n=26 cells from 3 independent transfections; ratio was calculated as F_478nm_/F_390nm_ for spectral configuration shown in **a**). (c) Binding titration curves for KRaION1 and KRaION2 presented with normalized excitation ratio F_478nm_/F_390nm_ (n=16 and 26 cells from 3 independent transfections for KRaION1 and KRaION2, respectively; data points, mean, error bars, standard deviation). Experiments were done at 33 °C.

Using the fluorescence ratio values measured from this calibration experiment, we calculated the sensors’ K_d_ values when expressed in HeLa cells as 23±3 (mM) for KRaION1 and 20±2 (mM) for KRaION2 (**Figure 4c**). These K_d_ values are lower than, and inconsistent with, the K_d_ values obtained in vitro when measured under isotonic conditions, above. Similar discrepancies in K_d_ were observed for previously published genetically encoded potassium indicators as well, and thus potentially all potassium indicators described to date could provide inaccurate potassium measurement readouts when used in living cells, at least when compared to in vitro data. When performing the same calibration experiment with GINKO1, for example, we obtained an intracellular K_d_ value of 3±0.23 (mM) -- quite different from its isotonic value.

We attempted to understand this K_d_ discrepancy by measuring the binding affinity of indicators at different ionic strengths in vitro. We chose to use buffers containing a constant 5 mM Na^+^ and variable K^+^ in the range of 0.1 mM to 250 mM as we attempted to mimic intracellular concentrations of Na^+^ and K^+^. When measured in these conditions, we obtained a different K_d_ value for KRaION1 of 42.19±9.84 (mM), lower than in the isotonic case, and closer to the HeLa cell case. Different K_d_ values were also observed in all measured indicators under these conditions, with most appearing lower than in the isotonic case (**Supplementary Table 5**). This suggests that the indicators are sensitive to changes in ionic strength when measured in vitro, and full characterization of the binding affinity of the indicators may need to be performed in the cell type and physiological context of interest. In particular, future potassium indicator engineering efforts may benefit from screens being fully performed in a cell type of interest rather than in vitro, as we have recently described in the context of robotic molecular screening for the case of infrared fluorescent proteins and fluorescent voltage indicators, but which has not yet been applied to potassium indicators (Piatkevich *et al*, 2018).

## Discussion

We have designed a set of intracellular genetically encoded potassium indicators based on the insertion of Ec-Kbp mutants and homolog variants into the mNeonGreen fluorescent protein. The insertion of Ec-Kbp into mNeonGreen results in an indicator that is bright, fast, and has an isotonically measured binding affinity for potassium of 69±10 (mM). We named this indicator KRaION1. We sought to increase this K_d_ value to numbers closer to the intracellular concentrations of K^+^ in mammalian cells. We used two approaches: structure-guided mutagenesis and identification of alternative potassium binding proteins. First, we utilized NMR to solve the structure of Ec-Kbp that allowed us to reveal the potassium binding pocket which coordinates the potassium ion through the backbone carbonyls of residues V7, A10, G75, I77, and I80. With this information, we rationally designed two mutants with higher isotonically measured K_d_ values, mNG-Ec-Kbp-D9N and KRaION2, with values of 138±21 (mM) and 96±9 (mM), respectively. In parallel to mutagenesis, we also identified four Ec-Kbp homologs that have a conserved binding site for potassium and can be used as alternative binding moieties when inserted into the genetically encoded indicator. These homologs had not previously been confirmed as potassium binding proteins. When expressed intracellularly, both KRaION1 and KRaION2 maintain their functionality. However, the K_d_ measured in the intracellular environment for both indicators was ∼20 (mM), which is lower than observed in vitro. As noted, measuring K_d_ in a context where sodium was held constant, gave values intermediate between the isotonic condition and the live cell condition, raising the point that physiological characterization of potassium indicators may need to occur in the exact cell type and physiological context of interest.

Proteins’ monovalent cation binding sites are notoriously difficult to detect by all of the techniques used for structural biology. In NMR spectroscopy, local chemical shift differences between the free and ligand-bound forms of proteins often provide powerful circumstantial evidence of the location of ligand binding sites but this approach was not feasible for Ec-Kbp due to the substantial global conformational change between the free and bound forms and the extreme line-broadening of many signals in its free form. More direct evidence for the location of potassium binding sites can be obtained using ^15^N labelled ammonium ions to substitute for K^+^ (Werbeck *et al*, 2014; Eichmann *et al*, 2019). However, Ec-Kbp unfolds at the low pHs at which the ammonium ^1^H signal is distinct from the water ^1^H signal. Here we exploited the favorable NMR characteristics of thallium (Gill *et al*, 2005), well known as a good potassium substitute, to make what we believe is the first reported observation of scalar (J) coupling between a protein’s ^13^C enriched carbonyl carbons and a bound thallium ion. The large J couplings observed provide direct and unambiguous evidence of which groups coordinate the ion. In fact, the variation in J for the different residues suggests that the data may contain additional information about the interatomic distances and angles that could be decoded in the future.

Our refined potassium-bound structure of Ec-Kbp shows that its potassium binding site is formed by a pair of closely opposed loops that form tight turns with the ion coordinated by five backbone carbonyls. Pentavalent coordination of potassium in proteins is commonly observed (Harding, 2002), but it is also possible that a water molecule could be involved in coordinating the ion raising the coordination number to an even more commonly observed number. The structure reveals a likely explanation for Ec-Kbp’s specificity for potassium and slightly larger ions over sodium: the two ion binding loops are already tightly packed with potassium bound, while accommodating sodium would dictate a shortening of each of of the ion-carbonyl distances by approximately 0.36 to maintain favorable interactions. Such a compaction of the structure does not appear to be sterically feasible. The structure also explains the roles of most of the conserved residues surrounding the ion binding site and explains why it is difficult to find mutations that subtly alter potassium affinity without effectively abolishing it altogether.

The solution structure of Ec-Kbp, which details information about the potassium binding site, allowed us to rationally design the potassium indicator to alter its binding affinity without disrupting potassium binding. Variants D9N and E12A (KRaION2), which both display higher isotonically measured K_d_ values than KRaION1, were also brighter indicators, with ΔF/F_max_ of 434% and 318%, respectively. KRaION2 in particular, exhibited faster association kinetics with a τ_fast_ of 26 ms, which is faster than KRaION1 and the previously developed GINKO1 indicator. Additionally, the indicators made with the newly identified Ec-Kbp homologs exhibit a wide range of isotonically characterized K_d_ values, with the lowest K_d_ at 39±7 (mM) for mNG-C-Kbp and the highest K_d_ at 112±35 (mM) for mNG-D-Kbp.

It is notable that KRaION1’s apparent K_d_ for potassium is already more than two orders of magnitude higher in vitro than Ec-Kbp’s. Perhaps Ec-Kbp’s ability to change conformation is frustrated by having its N- and C-termini tethered by the constraints on their anchor points imposed by mNG’s structure. This suggests that further optimization of the linker sequences to mNG may prove fruitful for tuning future KRaIONs’ useful potassium concentration response ranges.

Despite successfully adjusting the binding affinities when indicators were tested in vitro, the intracellular K_d_ value of the genetically encoded potassium indicators KRaION1 and KRaION2 was not consistent with what was observed in vitro under isotonic conditions. This discrepancy in K_d_ highlights two important factors to consider and that require further study in the potassium indicator field: first, the conditions in which in vitro characterization are performed are important, and second, further work is necessary to achieve proper calibration of potassium in living cells. When testing the indicators in constant sodium conditions rather than isotonic conditions, we observed smaller K_d_ values, which were nonetheless still mismatched to the K_d_ values obtained in living cells. These kinds of discrepancies in K_d_ have been observed and explored for small molecule calcium and potassium probes (O’Malley *et al*; Rana *et al*, 2019). For potassium probes, a combination of several ionophores along with sucrose (for keeping cell volume constant and improving cell viability) were found to be most effective in potassium calibration experiments in Jurkat and U937 cell types. Dequenching of the probes was also observed and attributed to protein binding. However, genetically encoded indicators may behave differently when expressed intracellularly and could experience unwanted interactions with other cellular components that are currently unknown to us. On this basis, we suggest that further work is necessary to improve potassium calibration experiments for genetically encoded indicators. A similar strategy from Rana, would be to identify the best combination of ionophores needed to properly equilibrate the intracellular and extracellular ion concentrations in the cell types of interest. If issues persist, it may be necessary to look into any unwanted interactions that could affect the indicator’s performance in cells.

Before KRaION1, KRaION2, and the other variants described are used in biological contexts, we would advise characterization of these genetically-encoded potassium indicators, and calibration of potassium concentration, in the cells and physiological contexts of interest. As for the genetically-encoded indicators themselves, structure-guided mutagenesis can be combined with other techniques such as high-throughput, multi-parameter directed evolution approaches (Piatkevich *et al*, 2018) to further improve other parameters of interest, such as brightness and kinetics, especially in the context of cell types of interest.

## Materials & Methods

### Molecular cloning and mutagenesis

*De novo* gene synthesis of the designed potassium sensor including Ec-Kbp and identified homologs and subsequent subcloning into the pBAD-HisD vector were done by Genscript. The genes were codon-optimized for human cells. Mutagenesis of individual amino acid residues in KRaION1 was done using the QuikChange Site-Directed Mutagenesis Kit (Agilent). Forward and reverse primers used to generate each mutant are provided in **Supplementary Table 6**. The products of each mutagenesis reaction were transformed into TOP10 (Invitrogen) electrocompetent *E. coli* cells and grown on LB plates with 1:1000 dilution of carbenicillin and 0.002% arabinose. Fluorescent colonies were identified and sent out for sequencing (Eton Bioscience Inc.) for confirmation of the correct mutation for each construct. For mammalian expression, the KRaION1 and KRaION2 genes were cloned into a mammalian expression vector pN1 (Clontech). The DNA sequence of GINKO1 was obtained from Addgene (plasmid #113112).

### Indicator expression and purification

Protein expression and purification was conducted by transforming sensor constructs in TOP10 (Invitrogen) electrocompetent or chemically competent *E. coli* cells. Sensor constructs were expressed under an arabinose promoter and cells were grown in minimum RM medium (1X M9 Salts, 2% Casamino Acids, 0.2% Glucose, 1 mM Magnesium Chloride, 1 mM Thiamine) supplemented with 1:1000 carbenicillin and 0.002% arabinose for 20-24 hours at 37°C and another 24 hours at 18°C. Cells were then pelleted by centrifugation at 4,500 rpm for 20 minutes and stored at –20°C. On the day of purification, cells were thawed at room temperature and were re-suspended in 1X PBS, 300 mM NaCl with 50X Lysozyme (100 mg/mL). Cells were kept on ice and were sonicated for 20 minutes at 20% intensity. After sonication, samples were centrifuged at 5,000 rpm for 1 hour to pellet cell debris. The resulting supernatants were transferred to clean conical tubes for binding to the purification resin. Sample supernatants were incubated with 4 mL of Ni-NTA Agarose (Qiagen) for >1 hour at 4°C. After incubation, the beads were washed 3 times with 1X PBS, 300 mM NaCl solution and packed into a gravity flow column. The proteins were eluted with 1X PBS, 300 mM NaCl, 100 mM EDTA (pH 7.6) solution and dialyzed against various buffers depending on the subsequent measurement conditions (see below sections), and stored at 4°C.

### In vitro characterization of KRaION1 in non-isotonic conditions

To characterize KRaION1 in vitro, the following buffers were prepared: 150mM NaCl, 25mM HEPES, 25mM MES and 10mM Tris (to be used as 0 mM K^+^) and 150mM KCl in 25mM HEPES, 25mM MES and 10mM Tris. Absorbance was measured at 250–600 nm range at both 0 mM K^+^ and 150 mM K^+^ using a spectrophotometer (Hitachi SHIMADZU UV-3600 Plus). The extinction coefficients were determined using alkine denaturation with 1M NaOH as described previously (Piatkevich *et al*, 2010). Fluorescence was measured using two excitation wavelengths at 400 nm and 480 nm and emission ranges of 410–700 nm and 490–700 nm, respectively, in the same buffers with 0 mM K^+^ and 150 mM K^+^. Quantum yields were measured at 0 mM and 150 mM K^+^ buffers at excitation wavelengths 405 nm and 480 nm. Fluorescence spectra and quantum yields were measured using the Edinburgh FLS1000 spectrometer.

For pK_a_ measurements, 0 mM K^+^ and 150 mM K^+^ buffers were prepared at varying pH levels: 2.6, 3.0, 4.0, 5.0, 7.0, 7.5, 8.0, 9.0. The pH of the buffers was adjusted by using HCl or NaOH. Fluorescence response of the indicators during this pK_a_ titration was measured at a range of 250–600 nm. pKa measurements were done using Plate Reader Thermo Varioskan LUX. Stopped flow spectrophotometry measurements to measure kinetics of the sensors were obtained at an excitation wavelength of 480 nm and collected at 520 nm at 40 mM K^+^ concentration using Applied Photophysics Ltd, SX 20.

For K^+^ titration, purified proteins in artificial mammalian cell cytoplasm buffer (12 mM NaHCO_3_ and 1 mM MgCl_2_ in 25mM Tris/MES, pH 7.4) were diluted into a series of buffers with K^+^ (potassium D-gluconate) concentration ranging from 0 to 260 mM. For examining K^+^ specificity, purified proteins were diluted into a series of buffers that each contained a different salt, these being: LiCl, NaCl, KCl, RbCl, CsCl, NH_4_Cl, and MgCl_2_ ranging from 0 to 260 mM. The fluorescence spectrum of the purified proteins in each solution (150 µL) was measured using a Thermo Scientific Varioskan LUX microplate reader with excitation at 490 nm and emission from 510 nm to 600 nm.

### In vitro characterization of binding affinity (K_d_) in isotonic conditions

Protein expression and purification was done as previously described. For K_d_ characterization experiments, we utilized Amicon Ultra-4 10K Centrifugal Filter units to exchange elution buffer for characterization buffer, which was 25 mM MES, 25 mM HEPES, pH 7.3. After buffer exchange, sensors were added to a 96-microwell plate, where each well had an isotonic solution that varied concentrations of KCl to NaCl ratio. KCl concentrations tested were in the range of 0 mM to 700 mM KCl. For characterization of K_d_ in a secondary condition, we utilized 25 mM MES, 25 mM HEPES, pH 7.3 to make solutions with a constant 5 mM NaCl concentration and increasing concentrations of KCl in the range of 0.1 mM to 250 mM K^+^. Absorbance and fluorescence measurements were taken with a Tecan Plate Reader (Tecan Spark) at temperature range 24 - 26°C. For each characterized indicator, we measured from three technical replicates.

Titration curves and K_d_ were later obtained by fitting into the following equation using a custom Python script: *Q* = (*Q*_*max*_ - *Q*_0_)*Y* + *Q*_0_ where Q is fluorescence and Q_max_ and Q_0_ are maximum and minimum fluorescent yields, respectively. Y is 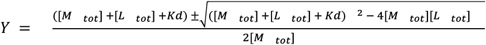 where M is a fixed concentration component of the genetically encoded sensor, L is the varying concentration component of K^+^ and K_d_ is the dissociation constant. All calculated values are expressed as K_d_ ± standard deviation.

### Ec-Kbp.Tl^+^ NMR and Ec-Kbp.K^+^ structure determination

To identify the amino acid groups that ligand the cation, a 0.25 mM sample of U-^13^C, U-^15^N-labelled Ec-Kbp prepared as described previously (Ashraf *et al*, 2016) was diluted in 20 mM sodium phosphate pH 7.2, 0.01% w/v sodium azide and 5% D_2_O with the addition of 1 mM thallium sulphate (to achieve 2 mM Tl^+^). From a 3D HNCO experiment recorded (1024×27×128 complex points for sweep widths of 9615×2127.7×2413.8 Hz in ^1^H, ^15^N and ^13^C, respectively) at 14.1 T and 298 K on a Bruker AVANCE IIIHD spectrometer equipped with a TCI cryoprobe and processed conventionally, the cross peaks that were split by an additional coupling the in the ^13^CO dimension were identified making use of the backbone resonance assignment of Ec-Kbp.Tl^+^ (B.O. Smith, personal communications) in CCPN analysis software (Vranken *et al*, 2005).

The refined structure of the Ec-Kbp.K^+^ was calculated using the same restraints as in the original structure determination (Ashraf *et al*, 2016), but with the addition of a potassium ion and 5 distance restraints of 2.75+/-0.1 Å between it and the five coordinating carbonyls (Harding, 2006). The parameters for the singly charged potassium ion were added to the topallhdg5.3.pro and parallhdg5.3.pro files in the ARIA2.3/CNS software with the epsilon and sigma values set to 0.62 kJ mol^-1^ and 3.0695 Å in the OPLX section used during refinement in explicit water (Mamatkulov & Schwierz, 2018). Eight rounds of structure calculations starting from randomised coordinates in Cartesian space were executed with 100 structures each. The 20 structures from the final iteration with the lowest restraint energy were refined using the ARIA2.3/CNS software in explicit water. The refined structures were analysed and figures prepared using PyMOL and Inkscape. The Ec-Kbp.K^+^ structure has been deposited at the protein data bank (PDB:7PVC).

### Metagenomic search for alternative Ec-Kbp domains using protein BLASTp

To search for alternative potassium binding domains that could replace Ec-Kbp in the indicator, we used NCBI’s BLASTp protein to protein sequence alignment tool. The Ec-Kbp amino acid sequence (NCBI reference sequence: WP_000522415.1) was used to search against the metagenomic database (env_nr). No other parameters were set to perform the search. The obtained results were filtered by percentage similarity of the amino acid sequences and were picked manually. The E-value metric was also taken into consideration, where numbers closest to 0 would be identified as good matches.

### In vitro screening of KRaION1 mutants and homologs

Protein expression and purification was done as previously described. To screen mutants and homologs for binding affinity, we first utilized Amicon Ultra-4 10K Centrifugal Filter units to exchange protein elution buffer for characterization buffer, which was 100 mM Tris, pH 7.3. After buffer exchange, sensors were added to a 96-microwell plate, in which sensors were tested at different KCl concentration increments ranging from 0 mM to 310 mM KCl in 100 mM Tris, pH 7.3. Absorbance and fluorescence measurements were taken with a Tecan Plate Reader (Tecan Spark). Sensor titration curves were fitted and K_d_ values were obtained fitting into the following equation using a custom Python script: *Q* = (*Q*_*max*_ - *Q*_0_)*Y* + *Q*_0_ where Q is fluorescence and Q_max_ and Q_0_ are maximum and minimum fluorescent yields, respectively. Y is 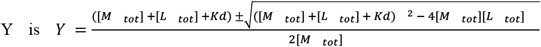 where M is a fixed concentration component of the genetically encoded sensor, L is the varying concentration component of K^+^ and K_d_ is the dissociation constant. All calculated values are expressed as K_d_ ± standard deviation.

### In vivo characterization and imaging in HeLa cells

HeLa cells (ATCC) were cultured in Dulbecco’s modified Eagle’s medium (Gibco) with 10% fetal bovine serum (FBS, YEASEN Biotech), and were incubated at 37°C with 5% CO_2_. Cells were plated on 12 mm coverslips (Fisher Scientific) coated with Matrigel in 24-well plates before transfection. Liposomal transfection method was applied according to the manufacturer’s protocol (YEASEN Biotech). HeLa cells were transfected by incubating with a mixture of 0.5 μg DNA and 1 μL Hieff Trans™ at room-temperature for 20 minutes. At 36-48 hours post-transfection, imaging was performed with an inverted wide-field Nikon Eclipse Ti2 microscope equipped with a SPECTRA III light engine (Lumencor), and a Orca Flash4.0v3 camera (Hamamatsu), controlled by NIS-Elements AR software, and using 20 × NA 0.75 objective lense..

The solutions with varying concentration of K^+^ were prepared by mixing two stock solutions (25 mM MES, 25 mM HEPES and 250 mM NaCl as the “no potassium solution” and 25 mM MES, 25 mM HEPES and 250 mM KCl with potassium) in the corresponding ratios. Cell medium was replaced with 500 μL of 25 mM, 25 mM HEPES and 250 mM NaCl buffer with 15 μM valinomycin and incubated for 15 minutes before perfusing. The solutions ranging from 0 mM to 150 mM K^+^ were administered with valinomycin (Aladdin Biochemical Technology Co., Ltd) at the final concentration of 15 μM right before perfusion.

To perform automated buffer exchange, the coverslip with cells was transferred to the RC-26G flow chamber (Warner Instruments, USA) and connected to a custom-built perfusion system consisting of peristaltic pump (Baoding Chuangrui Precision Pump Co., Ltd), the SV06 12 port switch value (Runze Fluid, China), controlled heated platform (Warner Instruments, USA), and vacuum pump. The system was designed to offer programmable buffer exchange and continuous monitoring of flow rate, temperature, and channel switching. The perfusion system was controlled by the custom LabVIEW code (National Instruments Corporation) and the experimental workflow was set up as illustrated:

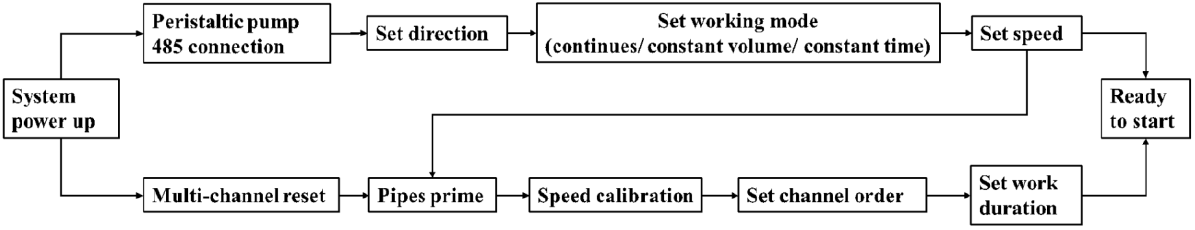

For live cell imaging, the programmable perfusion system was used to provide the series of extracellular buffers at 1 mL/min flow rate with different potassium concentration (0 mM to 150 mM K^+^ as described) containing valinomycin, ensuring that the cells were bathed in a consecutive environment without pipetting desired solution manually. The buffer temperature was kept at 33°C throughout the entire imaging.

## Data Availability

Sequences of the reported proteins are available at Genbank at the following accession codes: KRaION1, OK187805; mNG-Ec-Kbp-D9N, OK187806; KRaION2, OK187807; mNG-C-Kbp, OK187808; mNG-Pa-Kbp, OK187809; mNG-Hv-Kbp, OK187810; mNG-D-Kbp, OK187811. The solution structure of potassium bound Ec-Kbp has been deposited in the Protein Data Bank, accession code: <u>7PVC</u>. Plasmids of interest will be submitted to Addgene upon publication.

## Acknowledgments

We thank Orhan T. Celiker for help with Python script debugging. We are grateful to Yong Qian, Demian Park, Monique Kauke, and Ishan Gupta for useful discussions. We thank Dr. Zhong Chen from Instrumentation and Service Center for Molecular Sciences at Westlake University for the help with spectroscopy measurements. We thank Donald Campbell and Jared Williamson for assistance with preparing Ec-Kbp for NMR spectroscopy. We thank Chenlei Gu from Westlake University for helping with the development of the custom fluidic device. This work was supported by start-up funding from the Foundation of Westlake University, National Natural Science Foundation of China grant 32050410298, 2020 BBRF Young Investigator Grant, and MRIC Funding 103536022023 to K.D.P., an internal grant of the National Research Center Kurchatov Institute 𝒩<u>o</u>1056 from 02.07.2020 to F.V.S., and by Lisa Yang, John Doerr, HHMI, and grants NIH 1R01MH123977, NIH R01DA029639, NIH R01MH122971, NIH R01NS113499, NIH RF1NS113287, NIH RF1DA049005, NSF 1848029, NIH 1R01DA045549, NIH 1R01MH114031, NSF Grant 1734870, NIH R43MH109332, NIH R01GM104948, to E.S.B. C.C.T.C. was supported by the NSF GRFP Fellowship and Alfred P. Sloan Foundation scholarship.

## Acknowledgment to Henrietta Lacks and family

We acknowledge Henrietta Lacks and her family for their immense contributions to scientific research.

The mammalian cell experiments in this study were conducted on HeLa cells. The HeLa cell line was established from the tumor cells of Henrietta Lacks without her knowledge or consent. Since 1951, the use of the HeLa cell line has made countless contributions to medicine and science. E.S.B. makes a donation, on behalf of the lab, to the Henrietta Lacks Foundation for each publication the Boyden lab makes that uses HeLa cells, as inspired by the Reck-Peterson lab’s example.

## Author Contributions

C.C.T.C., K.D.P. and E.S.B. initiated the project. C.C.T.C., F.V.S., and K.D.P. designed the mNG-Ec-Kbp (KRaION1) indicator. C.C.T.C. and K.D.P. developed the indicators, performed mutagenesis, screening, and identified Ec-Kbp homologs. C.C.T.C. and M.Y. performed in vitro characterization and data analysis.

C.L. and K.D.P. performed sensor characterization in HeLa cells. L.Y. and K.D.P. performed ion selectivity experiments. B.O.S. determined the Ec-Kbp NMR structure and provided guidance on mutagenesis and data analysis. C.C.T.C., B.O.S., K.D.P. and E.S.B. interpreted the data and wrote the manuscript with contributions of all authors. K.D.P. and E.S.B. oversaw all aspects of the project.

