## Supplementary Information for "Tuning the sensitivity of genetically encoded fluorescent potassium indicators through structure-guided and genome mining strategies"

| | |
|---|---|
| **Supplementary Table 1** | NMR and refinement statistics for Ec-Kbp.$K^+$. |
| **Supplementary Table 2** | Physical properties of the tested KRaION1 mutants and homologs under screening conditions in solution. |
| **Supplementary Table 3** | Search results obtained from protein BLAST of the Ec-Kbp sequence. |
| **Supplementary Table 4** | Binding affinity and dynamic range characterization of genetically encoded indicators made with alternative binding domains. |
| **Supplementary Table 5** | Binding affinity ($K_d$) measurements obtained at varied conditions in vitro. |
| **Supplementary Table 6** | Primer sequences used for site-directed mutagenesis of Ec-Kbp. |
| **Supplementary Figure 1** | Structure of Ec-Kbp.$K^+$ and mutation sites. |
| **Supplementary Figure 2** | Emission spectra of mNG-Ec-Kbp and variants upon $K^+$ administration. |

**Supplementary Table 1.** NMR and refinement statistics for Ec-Kbp.$K^+$.

| | **Ec-Kbp.$K^+$** | |
|---|---|---|
| **NMR distance and dihedral constraints** | | |
| Distance constraints | | |
| Total NOE | 5626 | |
| Ambiguous | 2741 | |
| Unambigous | 2885 | |
| Intra-residue | 1237 | |
| Inter-residue | | |
| Sequential ($\|i – j\| = 1$) | 715 | |
| Medium-range ($\|i – j\| < 4$) | 371 | |
| Long-range ($\|i – j\| > 5$) | 562 | |
| CO to $K^+$ restraints | 5 | |
| Dihedral angle restraints[a] | | |
| φ | 81 | |
| ψ | 81 | |
| **Structure statistics** | | |
| Violations (mean and s.d.) | | |
| Distance constraints (Å) | $4.54\times10^{-2}\pm1.93\times10^{-3}$ | |
| Violations per structure > 0.5 Å | 1.5±0.87 | |
| Violations per structure > 0.3 Å | 15±3.4 | |
| Deviations from idealized geometry | | |
| Bond lengths (Å) | $5.63\times10^{-3}\pm1.75\times10^{-4}$ | |
| Bond angles (º) | $0.607\pm1.61\times10^{-2}$ | |
| Impropers (º) | $1.61\pm8.69\times10^{-2}$ | |
| Average pairwise r.m.s. deviation[b] (Å) | All residues | Ordered residues[c] |
| Heavy | 1.56±0.23 | 0.77±0.06 |
| Backbone | 1.02±0.22 | 0.43±0.06 |
| Ramachandran statistics (%) | | Ordered residues[d] |
| Favoured | | 93±2 |
| Additionally allowed | | 7±2 |
| Disallowed | | 0±0 |

[a]Chemical shift based, not used in refinement. [b]Pairwise r.m.s. deviation was calculated among 20 refined structures. [c]Residues 3-14, 25-50, 54-89, 91-101, 105-149. [d]Residues 3-14, 25-148.

**Supplementary Table 2.** Physical properties of the tested KRaION1 mutants and homologs under screening conditions in solution.

| Indicator | Absorbance (nm)[a] | Fitted $K_d$ (mM)[b] | $\Delta F/F_{max}$ (%)[c] | Brightness relative to KRaION1 (%)[d] | Folding efficiency (%)[d] |
|---|---|---|---|---|---|
| KRaION1 | 406/505 | 12±4 | 222 | 100 | 100 |
| mNG-Ec-Kbp-D9N | 405/504 | 27± 7 | 449 | 89 | 68 |
| mNG-Ec-Kbp-D9A | 405/504 | 17±5 | 506 | 46 | 25 |
| mNG-Ec-Kbp-E12A | 405/504 | 28±7 | 355 | 90 | 62 |
| mNG-Ec-Kbp-E12Q | 405/504 | 52±15 | 275 | 58 | 33 |
| mNG-Ec-Kbp-K8A | 405/505 | 17±15 | 164 | 91 | 72 |
| mNG-Pa-Kbp | 405/504 | 5±2 | 233 | 45 | 259 |
| mNG-C-Kbp | 405/505 | 19±6 | 204 | 104 | 126 |
| mNG-Hv-Kbp | 405/504 | 68±18 | 94 | 88 | 38 |
| mNG-D-Kbp | 405/504 | 7±3 | 173 | 37 | 26 |
| mNG-Ec-Kbp-A10G | 405/504 | n.d.[e] | n.d.[e] | 50 | 22 |
| mNG-Ec-Kbp-N76D | 405/504 | n.d. | n.d. | 20 | 11 |
| mNG-Ec-Kbp-I77A | 405/505 | n.d. | n.d. | 74 | 42 |
| mNG-Ec-Kbp-I77G | 405/504 | n.d. | n.d. | 46 | 47 |
| mNG-Ec-Kbp-I77S | 405/504 | n.d. | n.d. | 59 | 33 |
| mNG-Ec-Kbp-I80A | 405/504 | n.d. | n.d. | 36 | 34 |
| mNG-Ec-Kbp-I80G | 405/504 | n.d. | n.d. | 17 | 40 |
| mNG-Ec-Kbp-I80S | 405/504 | n.d. | n.d. | 65 | 33 |
| mNG-Ec-Kbp-D9K | 405/504 | n.d. | n.d. | 71 | 36 |
| mNG-Ec-Kbp-E12K | 405/505 | n.d. | n.d. | 2 | 34 |

[a]Absorbance peaks obtained at 160 mM $K^+$. [b]$K^+$ titration was conducted in non-isotonic conditions in a range of 0 - 310 mM $K^+$. Equation used to fit data and obtain $K_d$ is: $Q = (Q_{max} - Q_0)Y + Q_0$. [c]Max fluorescence dynamic range ($\Delta F/F_{max}$) values represent fluorescence change at baseline of 0 mM to the highest fluorescence value obtained within the range of 0 - 310 mM $K^+$. [d]Brightness and folding efficiency were expressed as a percentage relative to KRaION1 (i.e. KRaION1 = 100%). [e]For values listed as n.d. = not determined; these indicators did not exhibit a fluorescence change response to $K^+$ titration.

**Supplementary Table 3.** Search results obtained from protein BLAST of the Ec-Kbp sequence.

| Query[a] | Homolog[b] | Percent Identity (%) | Alignment Length (aa) | Number of mismatches | Gap Opens | Query Start (aa) | Query End (aa) | Sequence Start (aa) | Sequence End (aa) | E-value | Bit score | % Positives |
|---|---|---|---|---|---|---|---|---|---|---|---|---|
| WP_000522415.1 | MNG82101.1 (C-Kbp) | 72 | 148 | 41 | 0 | 1 | 148 | 1 | 148 | 3.11E-83 | 230 | 85 |
| WP_000522415.1 | NP_253865.1 (Pa-Kbp) | 55 | 148 | 64 | 2 | 1 | 148 | 1 | 145 | 3.07E-58 | 167 | 74 |
| WP_000522415.1 | SUS08588.1 (D-Kbp) | 53 | 147 | 63 | 4 | 1 | 147 | 1 | 141 | 1.84E-47 | 139 | 67 |
| WP_000522415.1 | VAV91021.1 (Hv-Kbp) | 45 | 148 | 76 | 3 | 1 | 148 | 1 | 143 | 1.54E-38 | 117 | 61 |

[a]Query corresponds to the GenBank accession number of the Ec-Kbp sequence utilized for sequence alignment. [b]Accession numbers of the homologs found by the metagenomic protein BLAST search.

**Supplementary Table 4.** Binding affinity and dynamic range characterization of genetically encoded indicators made with alternative binding domains.

| **Indicator** | **Fitted $K_d$ (mM)[a]** | **$\Delta F/F_{max}$ (%)[b]** |
|---|---|---|
| mNG-C-Kbp | 39±7 | 213 |
| mNG-Pa-Kbp | 48±6 | 262 |
| mNG-Hv-Kbp | 63±15 | 106 |
| mNG-D-Kbp | 112±35 | 147 |

[a]$K^+$ titration was conducted in isotonic conditions in a range of 0.1 – 700 mM $K^+$. Equation used to fit data and obtain $K_d$ is: $Q = (Q_{max} - Q_0)Y + Q_0$. [b]Fluorescence dynamic range ($\Delta F/F_{max}$) values represent percent max fluorescence change within the range of 0.1 – 700 mM $K^+$.

**Supplementary Table 5.** Binding affinity ($K_d$) measurements obtained at varied conditions in vitro.

| | Condition 1[a] | Condition 2[b] |
|---|---|---|
| **Indicator** | **Fitted $K_d$ (mM)** | **Fitted $K_d$ (mM)** |
| KRaION1 | 69±10 | 42±10 |
| mNG-Ec-Kbp-D9N | 138±21 | 100±6 |
| KRaION2 | 96±9 | 66±9 |
| mNG-C-Kbp | 39±7 | 34±5 |
| mNG-Pa-Kbp | 48±6 | 42±14 |
| mNG-Hv-Kbp | 63±15 | 133±22[c] |
| mNG-D-Kbp | 112±35 | 22±14 |
| GINKO1 | 17±7 | 21±10 |

[a]$K^+$ titration was conducted in isotonic conditions in a range of 0.1 – 700 mM $K^+$. [b]$K^+$ titration was conducted at constant 5 mM $Na^+$ with an increasing concentration of 0.1 – 250 mM $K^+$. [c]$K^+$ titration was conducted at 5 mM $Na^+$ with an increasing concentration of 0.1 – 650 mM $K^+$ for indicator to reach a binding plateau. Equation used to fit all data in this table to obtain $K_d$ is: $Q = (Q_{max} - Q_0)Y + Q_0$.

**Supplementary Table 6.** Primer sequences used for site-directed mutagenesis of Ec-Kbp.

| Mutant[a] | Primer Sequence 5'- 3' |
|---|---|
| Ec-Kbp-Fw-K8A | CGG CCT GTT CAA CTT CGT GGC CGA CGC CGG CGA GAA GC |
| Ec-Kbp-Rv-K8A | GCT TCT CGC CGG CGT CGG CCA CGA AGT TGA ACA GGC CG |
| Ec-Kbp-Fw-D9N | CGG CCT GTT CAA CTT CGT GAA GAA CGC CGG CGA GAA GCT GTG |
| Ec-Kbp-Rv-D9N | CAC AGC TTC TCG CCG GCG TTC TTC ACG AAG TTG AAC AGG CCG |
| Ec-Kbp-Fw-D9K | CGG CCT GTT CAA CTT CGT GAA GAA GGC CGG CGA GAA GCT GTG |
| Ec-Kbp-Rv-D9K | CAC AGC TTC TCG CCG GCC TTC TTC ACG AAG TTG AAC AGG CCG |
| Ec-Kbp-Fw-D9A | CGG CCT GTT CAA CTT CGT GAA GGC CGC CGG CGA GAA GCT GTG |
| Ec-Kbp-Rv-D9A | CAC AGC TTC TCG CCG GCG GCC TTC ACG AAG TTG AAC AGG CCG |
| Ec-Kbp-Fw-E12K | CGT GAA GGA CGC CGG CAA GAA GCT GTG GGA TGC CGT G |
| Ec-Kbp-Rv-E12K | CAC GGC ATC CCA CAG CTT CTT GCC GGC GTC CTT CAC G |
| Ec-Kbp-Fw-E12A | CGT GAA GGA CGC CGG CGC CAA GCT GTG GGA TGC CGT G |
| Ec-Kbp-Rv-E12A | CAC GGC ATC CCA CAG CTT GGC GCC GGC GTC CTT CAC G |
| Ec-Kbp-Fw-E12Q | CGT GAA GGA CGC CGG CCA GAA GCT GTG GGA TGC CGT G |
| Ec-Kbp-Rv-E12Q | CAC GGC ATC CCA CAG CTT CTG GCC GGC GTC CTT CAC G |
| Ec-Kbp-Fw-A10G | GGC CTG TTC AAC TTC GTG AAG GAC GGC GGC GAG AAG CTG TGG GAT GC |
| Ec-Kbp-Rv-A10G | GCA TCC CAC AGC TTC TCG CCG CCG TCC TTC ACG AAG TTG AAC AGG CC |
| Ec-Kbp-Fw-N76D | CTG GTG GCC GTG GGC GAC ATC AGC GGA ATC GCC AGC |
| Ec-Kbp-Rv-N76D | GCT GGC GAT TCC GCT GAT GTC GCC CAC GGC CAC CAG |
| Ec-Kbp-Fw-I80A | CGT GGG CAA CAT CAG CGG AGC CGC CAG CGT GGA CGA TCA AG |
| Ec-Kbp-Rv-I80A | CTT GAT CGT CCA CGC TGG CGG CTC CGC TGA TGT TGC CCA CG |
| Ec-Kbp-Fw-I80G | CGT GGG CAA CAT CAG CGG AGG CGC CAG CGT GGA CGA TCA AG |
| Ec-Kbp-Rv-I80G | CTT GAT CGT CCA CGC TGG CGC CTC CGC TGA TGT TGC CCA CG |
| Ec-Kbp-Fw-I80S | CGT GGG CAA CAT CAG CGG AAG CGC CAG CGT GGA CGA TCA AG |
| Ec-Kbp-Rv-I80S | CTT GAT CGT CCA CGC TGG CGC TTC CGC TGA TGT TGC CCA CG |
| Ec-Kbp-Fw-I77A | GGT GGC CGT GGG CAA CGC CAG CGG AAT CGC CAG CG |
| Ec-Kbp-Rv-I77A | CGC TGG CGA TTC CGC TGG CGT TGC CCA CGG CCA CC |
| Ec-Kbp-Fw-I77G | GGT GGC CGT GGG CAA CGG CAG CGG AAT CGC CAG CG |
| Ec-Kbp-Rv-I77G | CGC TGG CGA TTC CGC TGC CGT TGC CCA CGG CCA CC |
| Ec-Kbp-Fw-I77S | GGT GGC CGT GGG CAA CAG CAG CGG AAT CGC CAG CG |
| Ec-Kbp-Rv-I77S | CGC TGG CGA TTC CGC TGC TGT TGC CCA CGG CCA CC |

[a]Each mutant has a forward (Fw) and reverse (Rv) primer sequence.

**Supplementary Figure 1.** Structure of Ec-Kbp.$K^+$ and mutation sites.

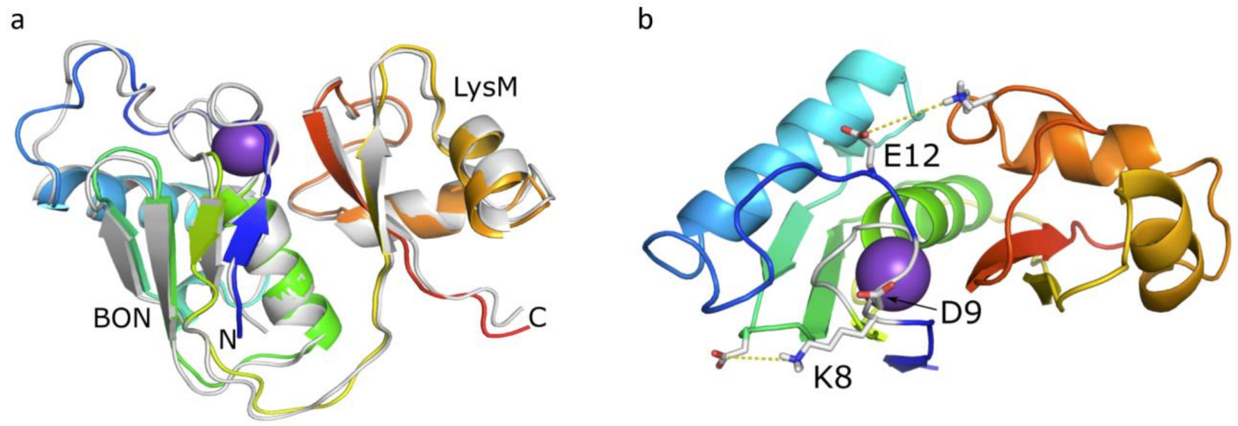

(a) Refined structure of Ec-Kbp in it's potassium bound state, shown as the rainbow-colored cartoon, is overlaid with the previously calculated structure of Ec-Kbp (PDB:5FIM) in grey. The potassium ion is represented by a purple sphere. (b) Location of the K8, D9, and E12 residues (shown as sticks) that were chosen for mutation in respect to the potassium binding site. The potassium ion is represented by a purple sphere and the protein backbone shown in cartoon representation colored from blue at the N-terminus through to red at the C-terminus. Residues D52 and K133, which may make ionic interactions with K8 and E12, respectively, are also shown in stick form with yellow dashed lines connecting them.

**Supplementary Figure 2.** Emission spectra of KRaION1 and variants upon $K^+$ administration.

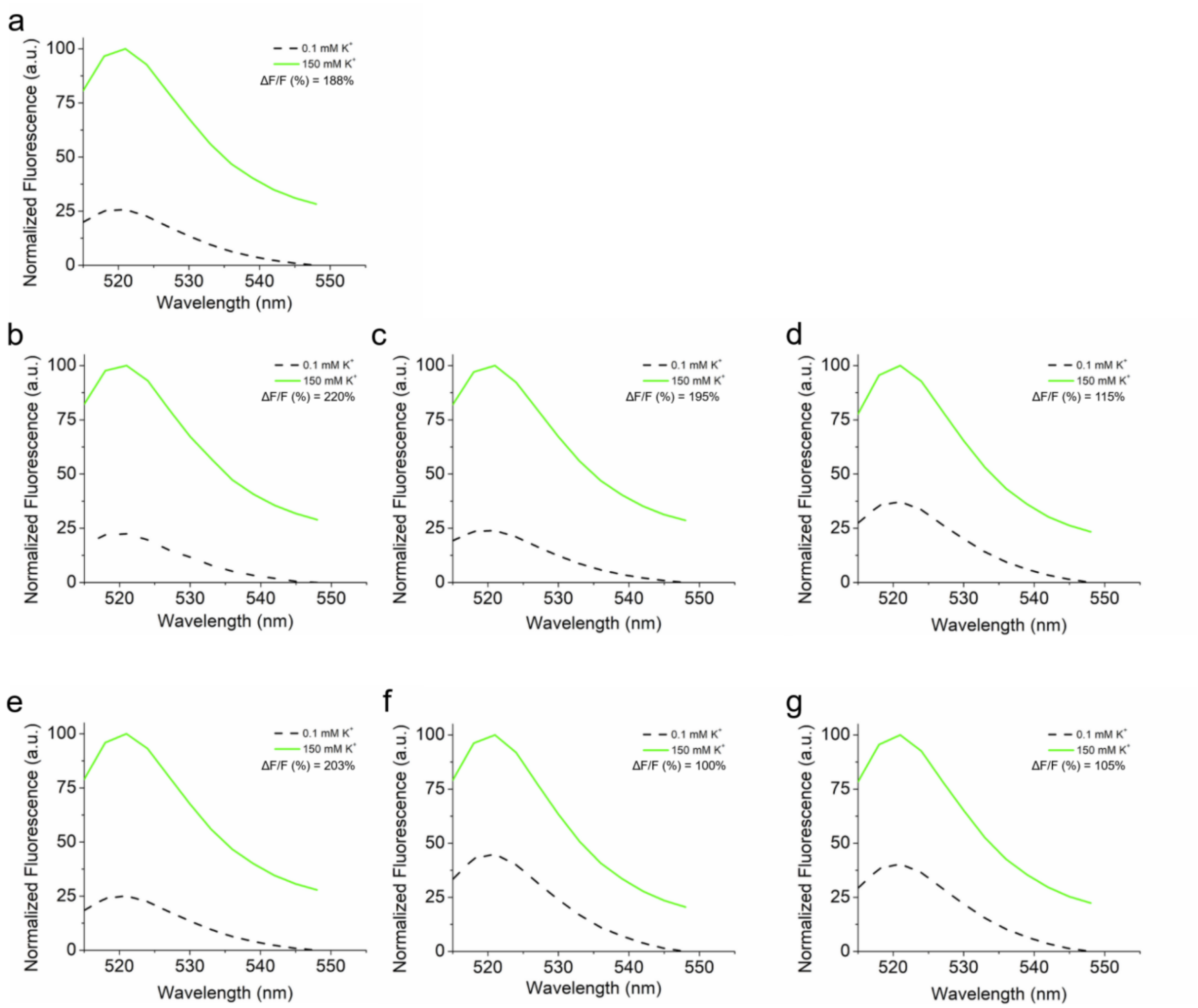

(a, b, c, d, e, f, g) Fluorescence emission spectra of KRaION1, mNG-Ec-Kbp-D9N, KRaION2, mNG-C-Kbp, mNG-Pa-Kbp, mNG-Hv-Kbp, and mNG-D-Kbp, respectively, at 0.1 mM and 150 mM potassium at pH = 7.3. ΔF/F refers to the maximum fluorescent change observed when measured in the concentration range of 0.1 - 150 mM $K^+$.
